# Global distribution and diversity of prevalent sewage water plasmidomes

**DOI:** 10.1101/2022.02.25.482059

**Authors:** Frederik Teudt, Saria Otani, Frank M. Aarestrup

## Abstract

Sewage water from around the world contains an abundance of short plasmids, a number of which harbor antimicrobial resistance genes (ARGs). The global dynamics of plasmid-derived antimicrobial resistance and functions is only starting to be unveiled. Here, we utilized a previously created dataset of 159,332 assumed small plasmids from 24 different globally collected sewage samples. We investigated the detailed phylogeny as well as the interplay between their protein domains, ARGs, and predicted bacterial host genera to help understand the global sewage plasmidome dynamics.

A total of 58,429 circular elements carried genes encoding for plasmid-related features, and MASH distance analyses showed a very high degree of diversity. A single very diverse cluster of 520 predicted *Acinetobacter* plasmids was predominant among the European sewage water.

Based on functional domain network analysis, we identified three groups of plasmids, mainly replication and mobilization domains. However, these backbone domains were not exclusive to any given group. *Acinetobacter* was the dominant host genus among theta-replicating plasmids at these size ranges. They contained a reservoir of the macrolide resistance gene pair msr(E) and mph(E). Macrolide resistance genes were the most common resistance genes in sewage plasmidomes and found in the largest number of unique plasmids.

While msr(E) and mph(E) were limited to *Acinetobacter*, erm(B) was disseminated among a range of *Firmicutes* plasmids, including *Staphylococcus* and *Streptococcus*, highlighting a potential reservoir of antibiotics resistance for these pathogens from around the globe.

**Importance:** Antimicrobial resistance is a global threat to human health inhibiting our ability to treat infectious diseases. This study utilizes sewage water plasmidomes to identify plasmid-derived features, and highlights antimicrobial resistance genes, particularly macrolide resistance to be abundant in sewage water plasmidomes in *Firmicutes* and *Acinetobacter* hosts. The emergence of macrolide resistance in these bacteria hints that macrolide selective pressure exists in sewage water, and that the resident bacteria readily can acquire macrolide resistance via small plasmids.

## Introduction

The effectiveness of antibiotic treatment is under threat from the oncoming resistance epidemic, making antimicrobial resistance (AMR) a major problem for global health^1^.

The AMR epidemic is partly caused by the transmission of antimicrobial resistance genes (ARGs) between different bacterial clones and species^2,3^. One of the most common and easily identifiable type of ARG-carrying mobile genetic elements is plasmids^4^. Plasmids often contain beneficial genes that the host microbe can use to accessorize its genome; these genes allow the microbe to adapt to a variety of factors, such as the presence of metals, antibiotics and biosynthetic and carbohydrate functions^5,6^.

Plasmids also carry genes necessary for initiating their own replication. There are three types of replication among plasmids: Rolling circle replication, theta replication, and strand displacement replication^7^. Rolling circle replication relies on a replication initiator to nick one of the plasmid strands creating a free strand end which is then elongated^8^. Plasmid theta replication and strand displacement replication is more similar to chromosomal replication: The initiator protein and/or transcription of a preprimer causes the DNA to melt at replication origins, and regular host replication forks extend RNA primers^7^.

Plasmids also often carry mobilization genes, allowing them to be transferred from cell to cell through mating pores. Mobilization is similar to rolling circle replication with the crucial difference that the nicked strand is not immediately replicated; instead it is transferred through a mating pore to an acceptor cell. Most mobilizable plasmids do not encode the gene necessary for mating pore formation. Instead, they rely on the mating pores provided by other plasmids. Plasmids encoding a complete mating pore are called conjugative, while those that rely on the mating pores of others are called mobilizable^9^.

Studies have shown an extremely high variability of plasmids within and between reservoirs^10,11^. An example of the former is the plasmid-encoded component of the pangenome of *E. coli*: there is far greater variation in this component than in the chromosome-encoded component of the pangenome. Additionally, the plasmid variation is not explained by pure phylogeny, but niche-phylogeny interaction^10^. A concrete example of how niche shapes plasmid gene content has been shown in the family *Lactobacillaceae*: Plasmids of bacteria adapted to living in vertebrates contain predominantly anaerobic metabolic genes, while plasmids from free-living bacteria contain aerobic metabolic genes; the availability of oxygen thus clearly affects the plasmid gene content^11^. It also appears that some plasmids are transmitting ARGs globally, as seen with the recent emergence of the colistin resistance gene mcr-9. This gene is found on similar IncHI2-ST1 plasmids in Europe, North America and China, suggesting a common ancestry.^12^ Thus, studying and understanding the global diversity and similarity of plasmids could provide novel insight into the genomic epidemiology of microbial genes, including ARGs.

Obtaining comparable samples from global sources is however, a challenge. We have recently suggested urban sewage as a comparable matrix and used that successfully to study the global occurrence of antimicrobial resistance, the virome, and human populations^13–15^. We recently applied long read sequencing to explore the occurrence and diversity of the small plasmidome in 24 globally obtained sewage samples and assembled 165,302 contigs (159,322 circular), of which 58,429 carried genes encoding for plasmid-related and 11,222 for virus/phage-related proteins^16^. These plasmidomes were compared to conventional whole community metagenomic extractions from the corresponding samples, elucidating which AMR classes was preferentially found in the plasmidome of sewage communities. One of the highlights of this article were the abundance of macrolide resistance genes in the plasmidomes, e.g. erm(B) genes, mph(E), mef(A), and msr(D).

Here, we investigate these data further with the aim to find gene clusters by looking for conserved regions at domain level. We also apply machine-learning tools for linking the sequences to their most likely host and investigate the interplay between AMR, host, and backbone proteins.

## Material and Methods

### Single read assembly

Sewage water circular elements and their generation methods have been described in a previous publication^16^. Briefly, DNA was extracted from sewage samples from 24 samples from around the world (sample list in Supplemental Table 1), non-circular DNA was digested, and the circular DNA was amplified through rolling circle amplification. Libraries were prepared for both Oxford Nanopore and Illumina sequencing. Nanopore reads were trimmed, and short reads (<10 kb) were discarded. The raw reads consisted of tandem repeats from the rolling circle amplification step. These were assembled into the original circular sequence, which was then polished using Illumina reads with pilon^17^.

### PLSDB comparison

To compare our assemblies to known plasmids, we mapped them against the plasmid database PLSDB (version 2020_11_19^18^) using blastn (all NCBI-blast commands were done with version 2.11.0+^19^). Likewise, the PLSDB plasmid were BLASTed against the plasmid assemblies with NCBI-blast (this time, PLSDB as query, assemblies as subjects). If the PLSDB plasmid coverage and the assembly coverage were both above 90% for a given alignment, the pair was considered to be matching.

### Homology reduction

Single read assembly creates duplicates, as each read from a plasmid can give rise to a sequence in the assembly. The assembly from each sample was homology reduced using kma (1.3.0)^20^: Settings were: 80%. -k 16 -NI -Sparse - -ht 0.80 -hq 0.80 -and. Default k-mer size 16 was used and homology thresholds were set to 80%. Homology reduction was done independently for each sample. The low thresholds were chosen as a countermeasure to the high error rate of Oxford Nanopore sequencing.

### Phylogenetic Analysis

The longest 100 sequences from each of the 24 samples were individually sketched using MASH 2.2^21^ and a 2,400 by 2,400 distance table was calculated. Singletons (circular elements with distance 1 to all other circular elements) were removed from the table. A neighbour-joining tree was created using PHYLIP 3.697^22^ and visualized using Microreact^23^.

### Pfam prediction and protein classification

Circular elements were scanned for Pfam domains, these domain predictions have been used previously to classify elements as plasmids or phages^16^. Briefly, Prodigal (version 2.6.3)^24^ was used to predict proteins, which were scanned against the Pfam database (version 33)^25^ using hmmscan from HMMER (version 3.3.1)^26^.

Pfam domains were used classify the putative proteins on the circular elements (as in our previous publication, the e-value cutoff was 0.000001^16^. If multiple domains with acceptable e-values were predicted in a protein, the domain with the lowest e-value was chosen. Each protein was thereby represented by its most significant domain. The number of times each domain was assessed as the most significant domain on a protein was counted. The proteins of the Comprehensive Antibiotic Resistance Database (CARD)^27^ protein homolog model and protein overexpression model had Pfam domains predicted in the same manner. The Pfam domains found on CARD database proteins were added to our domain classification scheme as “Potential ARG”.

The combination of domains in each circular element was calculated. The amount of times a domain was found on each individual circular element was not considered. The number of times each combination was encountered in a sample was counted.

We attempted to remove phage contamination with PPR-Meta^28^ and SourceFinder^29^ (Supplemental Figure 1 and Supplemental Figure 2, respectively). However, neither method proved very useful. PPR-Meta falsely predicted almost half of the plasmid sequences to be phage-derived. SourceFinder predicted an equally large proportion of plasmids to be either chromosomal or phage-derived.

### Bipartite Network Graph

A bipartite network was made of the circular elements and the domains found on them. For the circular elements, a node was made for each unique domain combination in each sample. For example: Sixteen circular elements in the Indian sample only contained the domains Relaxase and MobC; they were represented as a single node. For the domain part, each Pfam found in the data was simply represented as a node. An edge was drawn between a domain and a circular element if the domain was found on the circular element. To continue the example, MobC and Relaxase were each represented as a node and the Indian node described above was connected to them both. Networks were visualized by Gephi (version 0.9.2)^30^. The size of nodes was estimated by their counts.

The color of the nodes were based on Pfam function. The classification scheme was adapted from previous publications^16,31^. The scheme was updated to include toxin-antitoxin systems as a separate group, and potential ARG domains found in the CARD databases.

### ARG detection

The macrolide resistance gene msr(E) contains an ABC transporter Pfam domain (ABC_tran), thus ABC_trans is marked as a potential ARG. However, not all ABC_trans-containing proteins are msr(E) or ARGs. To find *bona fide* ARGs, the homology-reduced circular elements were aligned against the ResFinder database^32^ with kma^20^ using the following parameters: bcNano on, bc=0.7, mem_mode on.

### Plasmid host prediction

The homology-reduced circular elements were analyzed with a random forest model plasmid host predictor created by Aytan-Aktug et al.^33^. The genus predicted with the highest probability was chosen as the host taxon.

## Results

### Length distribution of circular assemblies

Sewage water plasmids from 24 samples from 22 countries around the world (Supplemental Table 1), which have been described previously^16^, were compared to the PLSDB database. The circular elements (Figure 1 top panel) were predominantly small compared to the PLSDB plasmids.

**Figure 1:**
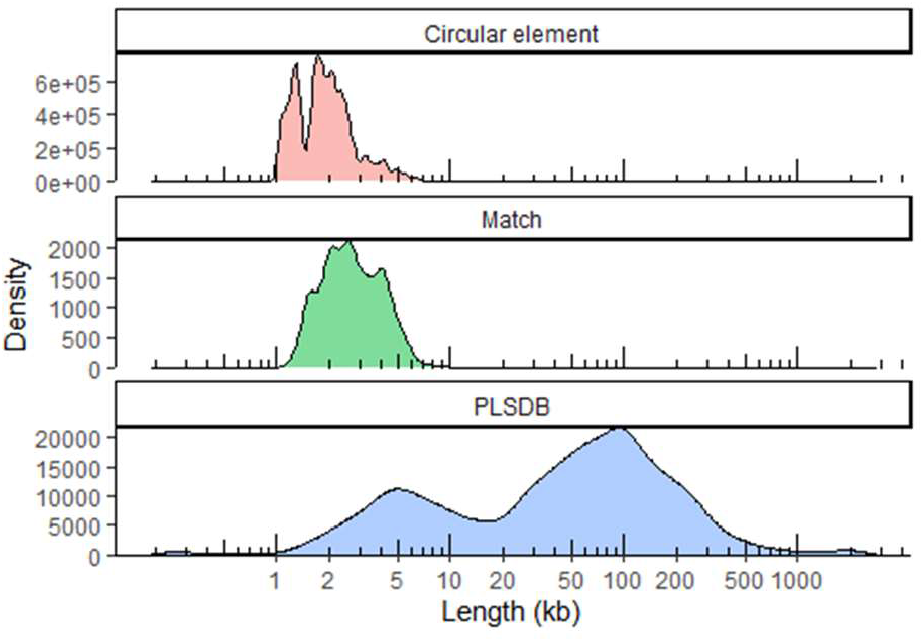
Length distributions of circular elements in our samples before homology reduction (top), PLSDB plasmids that match plasmids found in our samples (middle) and all PLSDB plasmids (bottom).

Plasmids from PLSDB^34^ were generally quite large (Figure 1 bottom panel). The PLSDB plasmids were split in two main types: Small plasmids of lengths of around 5 kb, and big plasmids ranging from 20 to 300 kb. The size range of PLSDB plasmids that matched circular elements in our samples were in the size range from about 1,500 bp to 5,000 bp (Figure 1 middle panel).

Overall, we are looking at a segment of plasmids that might be under-researched currently.

### Phylogenetic analyses

After individual homology-reduction of the 24 samples, the longest 100 circular elements from each sample were compared in phylogenetic analyses with neighbor-joining tree algorithm (Figure 2). With the exception of a smaller subset of predominantly European samples indicated by dotted lines, there was very little similarity between the analyzed circular elements (Figure 2). This was so despite the similar sizes of the tested circular elements (the largest subset from each sample).

**Figure 2:**
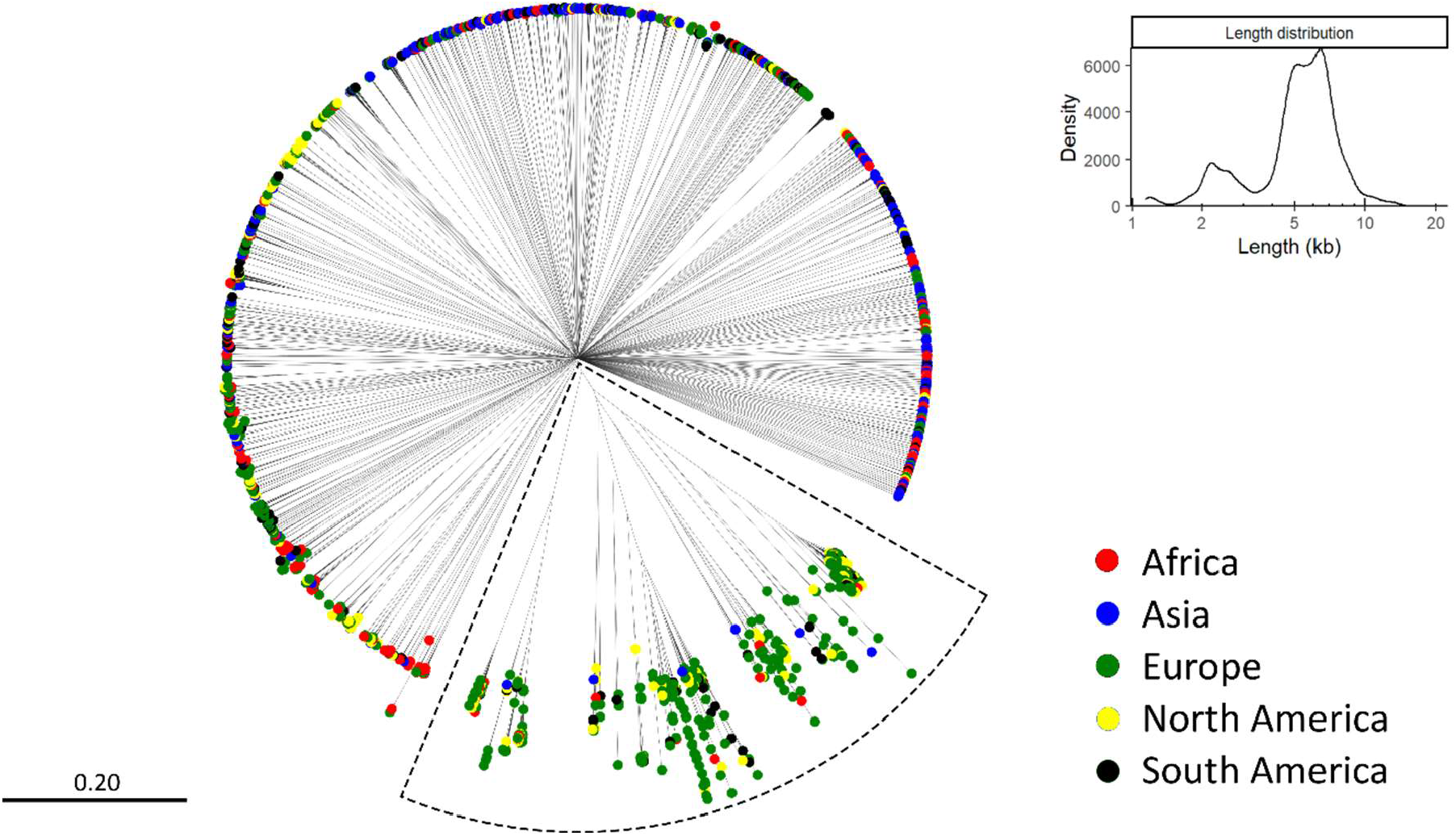
Dendrogram of the longest 100 circular elements from each sample. 140 singletons were removed leaving 2060 plasmids. Elements only connected through the center had no similarity. The length distribution of the circular elements is shown in the top right insert. The continent of origin of the plasmids is represented by their color.

The 24 samples were distributed across the continents as follows: four from Africa, five from Asia, eight from Europe, three from North America and four from South America. For this reason, overrepresentation European plasmids are expected. However, the cluster in the dotted box in Figure 2 has a higher proportion of European plasmids, than a third (which would match the proportion of samples that are from Europe). There are, however, African, Asian, North American, and South American plasmids in the cluster as well and this cluster was chosen for further downstream analyses.

### *Acinetobacter* plasmids from sewage water

The cluster of interest chosen from Figure 2 was dominated by plasmids predicted to be *Acinetobacter* with only a few *Neisseria* or *Bacillus* plasmids (Figure 3A). The plasmids in the cluster were still too many for visualizing their genes and alignments between them, so an even smaller subset was selected for visualization. The subset was composed of plasmids predicted to be *Acinetobacter* exclusively (Figure 3A dotted circle). A majority of the plasmids was European, but all continents were still present (Figure 3B). A plasmid from each branch of the subset was annotated and aligned against the other plasmids (Figure 3C).

**Figure 3:**
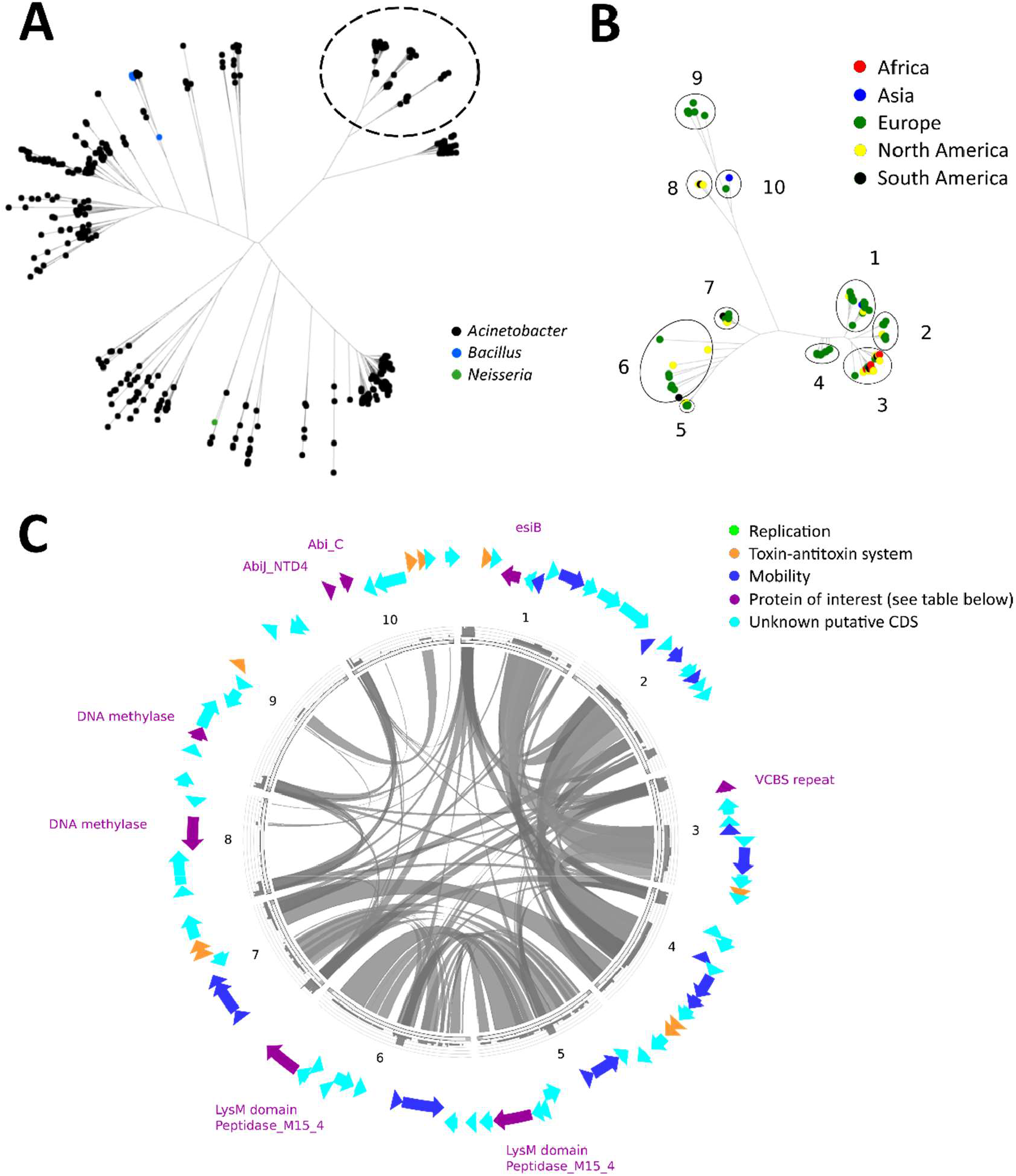
Visualizations of putative *Acinetobacter* plasmids. A: Dendrogram of the European cluster colored by genus. Subtree selected for closer inspection circled in black. B: Dendrogram showing geographical distribution of the selected *Acinetobacter* plasmids. Plasmids were divided into numbered groups from 1-10 for alignment-based comparison. C: Circos plots of representatives from each of the ten groups. Note that no replication genes were found. The functions of the genes highlighted in purple text is seen in Table 1.

This cluster was of interest because it contains no known replication mechanisms. This is despite the fact it contains other predominantly known plasmid-related functions, *i*.*e*., mobility systems and toxin-antitoxin systems^35,36^. The lack of replication proteins in these genetic elements indicates that this group of circular elements are either not plasmids, plasmids without replication initiator proteins, or plasmids with new types of replication initiator proteins.

Figure 3C shows the predicted ORFs in representatives from the subset, and Table 1 lists proteins of interest. Many of the ORFs did not have a predicted Pfam associated, or they were Pfams with very uncertain function, *e*.*g*., various helix-turn-helix domains. The proteins with assignable functions involved phage protection systems (N6_N4_Mtase^45^, N6_Mtase^46–48^, AbiJ_NTD4^49^, and Abi_C^50^); Cell wall metabolism (LysM^41,42^ and Peptidase_M15_4^43,44^); immune escape (esiB^39^); and cell-cell signaling (VCBS^40^).

**Table 1:**
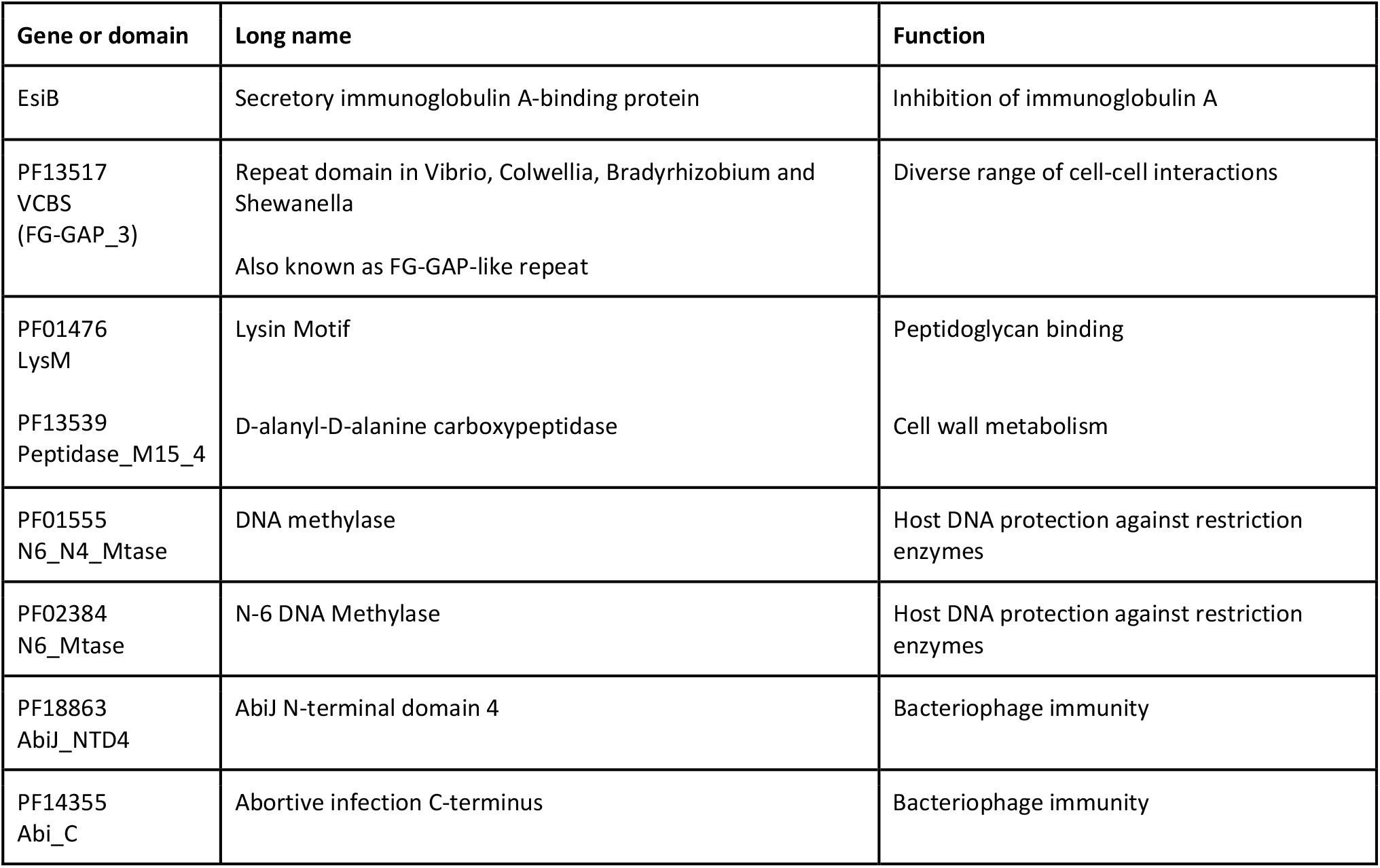
Protein families predicted on *Acinetobacter* cluster (purple CDS in Figure 3C).

| Gene or domain | Long name | Function |
| --- | --- | --- |
| EsiB | Secretory immunoglobulin A-binding protein | Inhibition of immunoglobulin A |
| PF13517<br>VCBS<br>(FG-GAP_3) | Repeat domain in *Vibrio*, *Colwellia*, *Bradyrhizobium* and *Shewanella*<br>Also known as FG-GAP-like repeat | Diverse range of cell-cell interactions |
| PF01476<br>LysM | Lysin Motif | Peptidoglycan binding |
| PF13539<br>Peptidase_M15_4 | D-alanyl-D-alanine carboxypeptidase | Cell wall metabolism |
| PF01555<br>N6_N4_Mtase | DNA methylase | Host DNA protection against restriction enzymes |
| PF02384<br>N6_Mtase | N-6 DNA Methylase | Host DNA protection against restriction enzymes |
| PF18863<br>AbiJ_NTD4 | AbiJ N-terminal domain 4 | Bacteriophage immunity |
| PF14355<br>Abi_C | Abortive infection C-terminus | Bacteriophage immunity |

### Clustering of circular elements into plasmids and bacteriophages

To investigate the circular elements at large, we made a network graph of all domains to trace their interactions in the global environment. The network layout was calculated by the ForceAtlas2^51^ algorithm. The layout revealed one main plasmid cluster and two clearly distinct bacteriophage clusters; the main cluster contained most of the nodes, while Phage clusters 1 and 2 were in the periphery (Figure 4). Interestingly, the two clusters were both apparently ssDNAs, as indicated by their domains^52–54^. One cluster contained viruses with replication domains Gemini_AL1 or Viral_Rep which may actually infect eukaryotes primarily^53^ (Figure 4 phage cluster 1); the other cluster contained no evident replication proteins, but did contain scaffolding protein Chlamy_scaf^52^ (Figure 4 phage cluster 2) and capsid domain Phage_F^54^ (pink), both found in ssDNA bacteriophages.

**Figure 4:**
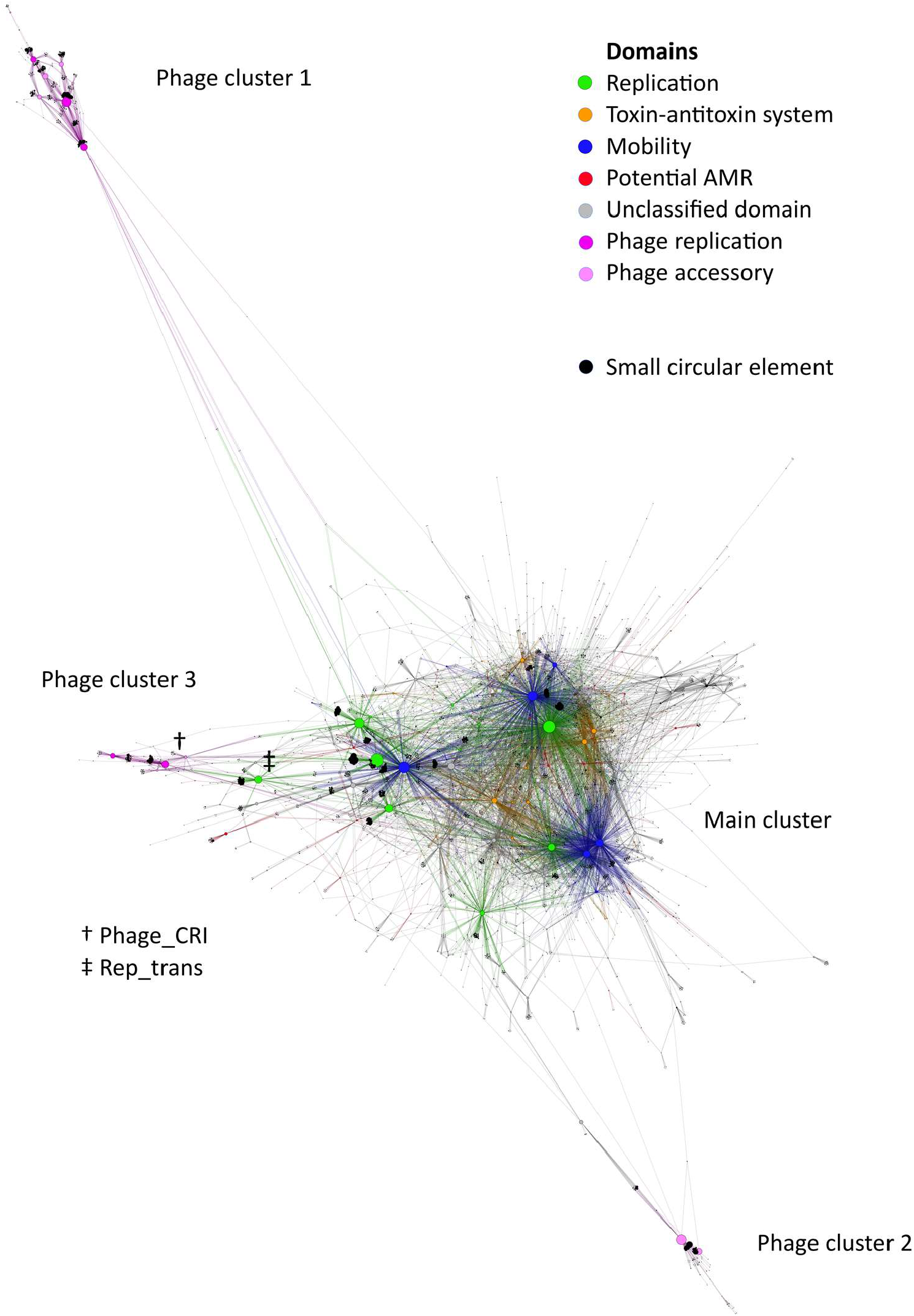
Bipartite network graph of sewage water Pfam domains. Layout by ForceAtlas2, node sizes ordered by count. The sequences separate into four distinct clusters. A main plasmid cluster, and a viral cluster at the top (phage cluster 1), at the bottom (phage cluster 2), and to the left (phage cluster 3, centered around the Phage_CRI domain). Phage cluster 3 was not well separated from the plasmids due to a stronger association to the replication initiator domain of the main cluster, most noticeably Rep_trans.

There is one last bacteriophage cluster, but it is less clearly separated from the main cluster (Figure 4 phage cluster 3). The majority of the viral accessory proteins of this cluster are either associated with the Phage_CRI or, to a lesser extent, the Phage_X viral replication domains found in this cluster. However, some are associated with the replication initiation domains of the main plasmid cluster, explaining why phage cluster 3 fails to separate from the main cluster. This is most noticeably from the fact that the Rep_trans domain has been pulled halfway from the main cluster towards phage cluster 3.

### Comparison of plasmid replication groups

Plasmid replication initiators were predictably the most common domains in the dataset (Figure 4, green nodes) with mobilization domains as close second (Figure 4, blue nodes). Replication and mobilization genes separate into three clusters in the network with 2-4 major domains in each. Group 1 is composed of Rep_1, Rep_2, RepL, and Mob_Pre and is the most prevalent. Group 2 is composed of Rep_3 and MobA_MobL. Group 3 is composed of Replicase, Relaxase, and MobC and is the least prevalent. There were more backbone proteins, but these were selected due to two factors. First, they needed to be large enough to be identifiable in the network as a major backbone domain. Second, they needed to be unambiguously clustered with other backbone in their vicinity. Rep_trans is closest to Group 1, but it is not included in the group. Of the 72,225 circular elements, 34,659 contained Pfam domains, and 21,639 had plasmid backbone proteins from at least one of the three replication groups.

Replication group 1 contained domains for rolling circle initiator proteins (Rep_1^55^, Rep_2^56,57^, RepL^58,59^, and the exluded Rep_trans^60^), while groups 2 and 3 contain domains of theta replication initiators (Rep_3^61,62^ and Replicase^63^). It therefore makes sense that group 1 replication domains are the ones that are the most entangled with bacteriophages, which also use rolling circle replication^8^. The rolling circle-replicating section of the network generally consisted of a few large nodes, while the theta-replicating section was more dispersed (Figure 5). Most of the major backbone domains had a large cluster of circular elements connected too them: the elements with only that single backbone domain found on them. These were the most common types of circular elements.

**Figure 5:**
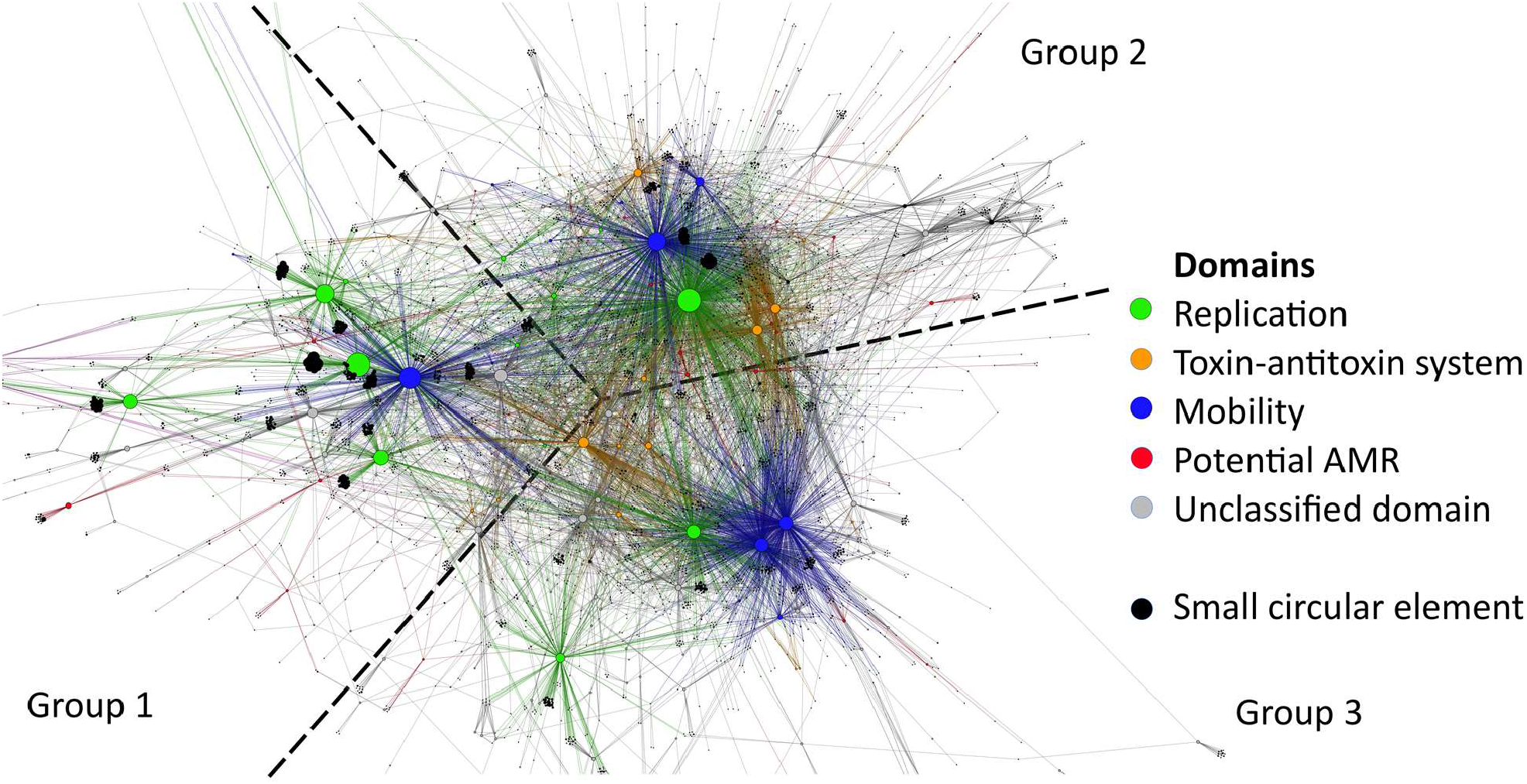
Plasmid region of bipartite network graph of sewage water Pfam domains. Layout by ForceAtlas2, node sizes by count. The backbone proteins (replication and mobilization, green and blue respectively) can be separated into three groups.

Closer inspection of the backbone genes showed that there are minor overlap between the backbone clusters (Table 2). As expected, not all replication proteins were associated with a mobilization protein. However, if replication proteins were associated with mobilization proteins, they would predominantly belong to specific Pfams. Two different replication proteins were almost never found on the same plasmid (Table 2). Likewise, different mobilization domains were generally not found together on the same plasmid with the exception of Relaxase and MobC which serve complimentary functions during plasmid mobilization (Table 2).

**Table 2:** Backbone types of plasmids. Numbers indicate the amount of times two domains were observed on the same plasmid. Rep_1, Rep_2, RepL, Rep_3, and Replicase are replication initiator domains. Mob_Pre, MobA_MobL, Relaxase, and MobC are mobilization-related domains.

|  |  | Group 1 |  |  |  | Group 2 |  | Group 3 |  |  |
| --- | --- | --- | --- | --- | --- | --- | --- | --- | --- | --- |
|  |  | Rep_1 | Rep_2 | RepL | Mob_Pre | Rep_3 | MobA_MobL | Replicase | Relaxase | MobC |
| Group 1 | Rep_1 | 7508 | 3 | 0 | 435 | 3 | 28 | 0 | 3 | 0 |
|  | Rep_2 | 3 | 2508 | 1 | 895 | 11 | 0 | 0 | 5 | 2 |
|  | RepL | 0 | 1 | 1418 | 401 | 0 | 8 | 0 | 19 | 19 |
|  | Mob_Pre | 435 | 895 | 401 | 4003 | 420 | 0 | 83 | 0 | 0 |
| Group 2 | Rep_3 | 3 | 11 | 0 | 420 | 5945 | 1457 | 4 | 175 | 173 |
|  | MobA_MobL | 28 | 0 | 8 | 0 | 1457 | 2386 | 101 | 0 | 0 |
| Group 3 | Replicase | 0 | 0 | 0 | 83 | 4 | 101 | 1063 | 344 | 381 |
|  | Relaxase | 3 | 5 | 19 | 0 | 175 | 0 | 344 | 1072 | 842 |
|  | MobC | 0 | 2 | 19 | 0 | 173 | 0 | 381 | 842 | 1118 |

The division of backbone genes into groups was based on the observed trend in the network but it was not a perfect division of the domains. Replication proteins of Group 2 and Group 3 most often mixed with mobilization domains of other groups, while Group 1 replication proteins predominantly associated with Mob_Pre.

Besides replication initiator proteins and mobilization proteins, the network also showed one other type of very common domains: toxin-antitoxin systems. They were more closely associated to the theta-replicating plasmids (Figure 5, orange nodes). Toxin-antitoxin systems were not as common as replication and mobility domains, but far more common than any other type of domains.

### ARGs and backbones

After grouping the backbone proteins into three categories, the ARGs associated with each type were investigated. Table 3 shows the backbone as they occur on plasmids that carried ARGs. Interestingly, many ARGs were found on Group 1 plasmids, but not those carrying Rep_1, despite it being the most prevalent replication protein. The most prevalent backbone domain on ARG-carrying plasmids was Mob_Pre (of Group 1 backbones). The next most prevalent protein was Rep_3 (of Group 2 backbones). Overall, it was clear that ARGs associated with specific backbone proteins, rather than all of a backbone group.

**Table 3:** Backbone types of ARG-carrying plasmids. Numbers indicate the amount of times two domains were observed on the same AMR-plasmid.

|  |  | Group 1 |  |  |  | Group 2 |  | Group 3 |  |  |
| --- | --- | --- | --- | --- | --- | --- | --- | --- | --- | --- |
|  |  | Rep_1 | Rep_2 | RepL | Mob_Pre | Rep_3 | MobA_MobL | Replicase | Relaxase | MobC |
| Group 1 | Rep_1 | 7 | 0 | 0 | 0 | 0 | 0 | 0 | 0 | 0 |
|  | Rep_2 | 0 | 27 | 0 | 21 | 2 | 0 | 0 | 0 | 0 |
|  | RepL | 0 | 0 | 2 | 0 | 0 | 0 | 0 | 0 | 0 |
|  | Mob_Pre | 0 | 21 | 0 | 39 | 4 | 0 | 0 | 0 | 0 |
| Group 2 | Rep_3 | 0 | 2 | 0 | 4 | 37 | 9 | 0 | 0 | 3 |
|  | MobA_MobL | 0 | 0 | 0 | 0 | 9 | 9 | 0 | 0 | 0 |
| Group 3 | Replicase | 0 | 0 | 0 | 0 | 0 | 0 | 2 | 2 | 2 |
|  | Relaxase | 0 | 0 | 0 | 0 | 0 | 0 | 2 | 9 | 5 |
|  | MobC | 0 | 0 | 0 | 0 | 3 | 0 | 2 | 5 | 8 |

Bacteriophages usually do not carry ARGs. To discover if that was the case in our samples, we investigated if there were any ResFinder hits found on the same circular elements that carried bacteriophage proteins. No bacteriophage domains were found alongside ResFinder hits.

Supplemental Figures 3 and 4 shows the location of the most commonly found ARG domains in the domain network with respect to predicted host genus and sample origin. The domains RrnaAD, Aminoglyc_resit and CAT were closer associated with the rolling circle plasmids, while APH MFS_1, and ABC_trans were found with the theta-replicating plasmids. Pentapeptide_4 was located on the outskirts of the plasmid clusters. Table 4 lists which commonly found ARGs these domains correspond to.

**Table 4:** Geographical distribution of the most common ARGs along with the name of their most significant Pfam domains.

| ARG | Africa | Asia | Europe | North America | South America | total | Pfam |
| --- | --- | --- | --- | --- | --- | --- | --- |
| erm(B) | 2 | 2 | 20 | 5 | 6 | 35 | RrnaAD |
| msr(E) | 3 | 0 | 17 | 0 | 0 | 20 | ABC_tran |
| mph(E) | 2 | 0 | 17 | 0 | 0 | 19 | APH |
| qnrB19 | 1 | 1 | 1 | 3 | 8 | 14 | Pentapeptide_4 |
| erm(T) | 4 | 0 | 3 | 1 | 0 | 8 | RrnaAD |
| lnu(A) | 0 | 2 | 3 | 0 | 1 | 6 | Aminoglyc_resit |
| tet(39) | 1 | 0 | 2 | 0 | 2 | 5 | MFS_1 |
| aph(3')-III | 0 | 0 | 0 | 0 | 3 | 3 | APH |
| cat | 0 | 0 | 3 | 0 | 0 | 3 | CAT |

### AMR and bacterial hosts

The host of the ARG-carrying plasmids were also predicted in the global sewage plasmidomes (Supplemental Table 3). We had a total of 150 ARGs found after homology-reduction. Using a plasmid host species predictor^33^, we investigated the host range of ARG-carrying plasmids. The ARG-carrying plasmids were limited to 14 different predicted genera.

Interestingly, the most common ARGs in the homology-reduced plasmids were capable of conferring macrolide resistance, but in different host ranges (Supplemental Table 3). The most commonly found ARG was erm(B). erm(B) is a methyltransferase that confers macrolide resistance by methylating rRNA^64^. It was found in five different genera of *Firmicutes*. The next most common genes were the gene pair msr(E) and mph(E). msr(E) is a ribosome protection protein, conferring resistance to a wide range of ribosome-targeting antibiotics, e.g. macrolides, lincosamides, and phenicols^65^. mph(E) is a macrolide phosphotransferase, conferring resistance to macrolides^66^. The other AMR gene found on more than two plasmids were qnrB19 in *Klebsiella* and *Salmonella* (which confers resistance to quinolones)^67^; erm(T) in the *Staphylococcus, Lactiplantibacillus*, and *Streptococcus*; lnu(A) in *Staphylococcus*, and *Lactiplantibacillus*; tet(39) in *Acinetobacter*; and lastly, cat and aph(3’)-III were both found in *Escherichia*.

### Genera and backbones

Having found a link between ARG and host bacterial genera, we then investigated the relationship between the bacterial host genus and plasmid backbone type. The host predictions are listed in Supplemental Table 2. The five most often predicted bacterial host genera were analyzed for their backbone gene content. Importantly, hosts were predicted for all circular elements, including for bacteriophages. Two different methods for excluding phages were tried, but they were both too inaccurate (Supplemental Figure 1 and Supplemental Figure 2). However, bacteriophages should not have plasmid backbone genes so their output will not skew the backbone composition analysis.

*Planococcus* was by far the most often predicted host (n=26,165), but relatively few backbone genes were found on these plasmids (Supplemental Table 4). Rep_2 and Rep_L made up an unusually large proportion of backbone genes of these plasmids. As *Planococcus* is predominantly a marine halophilic genus^68,69^ and relatively few of these circular elements carried plasmid backbone genes, it is likely that most of these predictions were false, likely stemming from phage contamination.

*Neisseria* was also predicted in high abundance (n= 11710, Supplemental Table 5). The abundances of the backbone genes were fairly similar to the abundances in general, though groups 2 and 3 were again slightly underrepresented. *Neisseria* is a common human and animal commensal^70^, thus it not unexpected to find it in sewage.

*Lactiplantibacillus* plasmids were very similar to *Neisseria* plasmids, but notably contained many backbone gene combinations across groupings (*e*.*g*., Replicase and Mob_Pr e, Supplemental Table 6). *Lactiplantibacillus* is found as a commensal^71^ and is used for silage production^72^, thus is very likely to be found in sewage water.

*Acinetobacter* plasmids were had the rest a high abundance of groups 2 and 3 backbones; in fact a majority of group 3 backbones were predicted to be from *Acinetobacter* (Compare table Table 2 and Supplemental Table 7). *Acinetobacter* is fairly ubiquitous and expected in sewage water^73^.

*Escherichia* plasmids had much fewer Rep_1 genes than the population in general, but they were otherwise fairly representative of the population (Supplemental Table 8). *Escherichia* is a common gut bacterium^74^, and thus likely present in sewage, lending credence to its prediction as a host. *E. coli* had by far the most varied range of backbone domains.

There was a clear link between genus and plasmid backbone genes but no genus was predicted exclusively to one group. Coloring the domain network according to predicted host genus also highlighted the association between backbone genes and host genus. Importantly, the theta-replicating plasmids were predominantly *Acinetobacter* plasmids while the rolling circle replicating plasmids were more varied in their host predictions (Supplemental Figure 3). As anticipated, the sequences predicted to be from Planococcus were not plasmids, but mainly viruses (Supplemental Figure 3, purple nodes).

### Geographical distribution of domains and ARGs

Coloring the domain network according to continent of origin showed that the theta-replicating plasmid cluster was dominated by European samples, while sequences in phage clusters 1 and 2 were predominantly from Asian samples (Supplemental Figure 4). The rolling circle plasmids were quite varied in regarding their continent of origin.

The European samples contained a disproportionate amount of the most common ARGs found in the data. erm(B) and qnr19 were the only genes detected on all continents. The gene pair msr(E) and mph(E) were found on many plasmids in Europe, only a few in African samples and no other continents.

The three resistance gene most often encountered in the homology-reduced dataset all conferred macrolide resistance. erm(B) was found in *Firmicutes* and mph(E) and msr(E) were found in *Acinetobacter*. Additionally, erm(B) was found across five different genera (Supplemental Table 3). erm(B) were found on all continents. European samples contained most erm(B) genes, followed by the Americas, and the least in Africa and Asia; Africa on the other hand contained the most erm(T) genes. mph(E) and msr(E) was predominantly found in Europe (Table 4).

The domains pentapeptide_4 (containing the gene qnrB19) and CAT (containing the gene cat) were not located within replication groups. qnr19 was found in *Salmonella* plasmids (Supplemental table 3); *Salmonella* plasmids was very rarely predicted in the data, but almost all qnr19 genes were located on them. qnr had a worldwide distribution (Table 4). cat was found on European samples (Table 4) in *Escherichichia* (Supplemental table 3). Plasmids predicted to be *Escherichia* is not rare in the data, but are not associated with either replication. Like CAT, *Escherichia* lies in between the groups in the domain network (Supplemental Figure 4).

## Discussion

### Assembly properties

In this study, we further analyzed, previously extracted and assembled short circular elements, mainly plasmids, from sewage water from around the world^16^. 48% of the circular elements encoded functional domains (Pfam); this low proportion was likely due to the low plasmid/chromosome DNA ratio and potential low copy numbers can make it difficult to detect plasmids or low quality of Oxford Nanopore reads which the plasmids were assembled from. Of the Pfam-containing elements, 62% contained plasmid replication initiator proteins or mobilization systems, suggesting that our findings are not artefacts.

The lack of longer sequences above 3,000 bp (Figure 1 top panel) could be caused by a disadvantage of assembling each plasmid from a single read; in this study reads numbers tend to drop with read length, so assembly is naturally limited at higher ranges as single reads are less likely to completely span longer plasmids. The smaller peaks in the distribution could be caused by the existence of a few high-abundance plasmids or phages. There were also minor peaks in the distribution of PLSDB plasmids found in our samples (Figure 1 middle), due to redundancy in PLSDB.

The fact we see several small elements compared to PLSDB is likely due to the presence of other non-plasmid circular elements, notably bacteriophages. It could also be due to a database bias caused by lack of research into plasmids that are too small to carry genes of interest (*e*.*g*. AMR genes). However, there was no way of telling if all these elements were artefacts or true plasmids or phages, though it is unlikely that the sequences we detected functional domains on were artefacts.

### Sewage *Acinetobacter* plasmids

*Acinetobacter* was a very important sewage water plasmid host in our findings; the only major cluster found in our phylogenetic analysis was composed of *Acinetobacter* plasmids (Figure 3A) and it was the dominant genus among Replication Groups 2 and 3 in the functional analyses(Supplemental Table 7 and Supplemental Figure 3). *Acinetobacter* plasmids also contained all the msr(E) and mph(E) macrolide resistance genes we found.

Of the subset of *Acinetobacter* plasmids that were chosen for closer analysis, a majority of them were from European samples (Figure 3B). With regards to functional domains, replication domains were the most dominant in all plasmids, however they were absent in this plasmid subset (Figure 3C). It could be that the assemblies are integrative conjugative elements (ICEs). These element can exist as circular ones and contain genes for conjugation similar to plasmids, but they do not contain replication gene^75^. Instead, they rely on integration into the host genome for replication. At odds with this theory is the lack of any predicted integrases and excisionases between the sequences. It could also be a case of missed replication genes as there are regions of homology with no predicted ORFs (Figure 3C). Shifting the start point of the sequence does not reveal any ORF in these areas. It should be noted that only a small amount of these sequences are homologous, thus it is possible that the lack of a replication genes in this cluster is simply a coincidence. Lastly, theta-replicating plasmids, which *Acinetobacter* plasmids often are (Supplemental Table 7), can initiate replication with only host factors^7^, which could also explain the missing replication genes.

It made sense we found phage protection functions on our *Acinetobacter* plasmids, as plasmid-encoded phage defense systems is a known strategy for combating phage infection^76,77^. We also found cell wall metabolism functions. Cell walls are common antibiotics targets, and the genes involved with cell wall metabolism are potential AMR genes. Specifically the Peptidase_M15_4 found on our subset (Figure 3 C and Table 1) is similar to VanY, a vancomycin resistance gene^43,44^. However, VanY exists in a much larger gene cluster than what could possibly fit on these plasmids. So if these genes somehow confer resistance to a cell wall-targeting antibiotic type, it may not be vancomycin, and it is not a homologous mechanism to VanY.

### Functional domains in sewage plasmidomes

While the findings from the inspected *Acinetobacter* subset were interesting, these plasmids were only a small fraction of the investigated circular elements. The domain network allowed us to investigate the plasmid population at large.

The analysis of the sewage plasmidomes at large was done based on functional Pfam predictions. This methodology intrinsically filtered out very small possible artefacts from the assemblies if they contained no Pfams. It also acted as a control for separating plasmids from phages using domain classification.

The Pfam network analysis showed that there were three predominant plasmid replication groups (Figure 5 and Table 2). In each group, there was an association between replication domain and mobilization domain. This could be because replication and mobilization proteins are dependent on the host machinery; thus, certain domains are favored by certain hosts, and are therefore found together. The bacterial host genus predictions backed up that different bacterial genera favor different backbone domains (Supplemental Figure 3). It was clear that there was an overrepresentation of specific combinations of genera and backbone domains. It did not appear to correlate with closely related bacterial taxa, as *Planococcus* plasmids (of phylum *Firmicutes*, Supplemental table 4) had similar backbones to *Neisseria* plasmids (of phylum *Proteobacteria*, Supplemental table 5). Backbone protein families thus do not seem to follow host phylogeny. The emergence of dominant backbone proteins within a genus must therefore have been a more recent event.

Replication proteins, mobility proteins, and toxin antitoxin (TA) systems were the most dominant proteins, respectively. Finding replication and mobility proteins was expected as they form the backbone of plasmids, however finding several TA systems was not expected. TA systems can be thought of as selfish genetic elements, securing the stability of the plasmid carrying it through post-segregation killing^78^. It could be speculated that the relative abundance of TA systems correlates to the small size distribution of our plasmids. The small size of the plasmids limits them to have barebones gene compositions. Replication genes, mobility genes and TA systems allow the plasmid to be inherited, to spread, and to secure their persistence in the host, respectively. Together these genes can be viewed as the minimal set of genes a plasmid needs to secure its survival. The small size of the plasmids studied entails that this minimal set of genes dominates the gene network (figure 5); if larger plasmids were under study, these would presumably contain a bigger proportion of genes outside of these three protein types. The role of TA systems as a third type of backbone proteins could explain why TA domains sometimes clustered together with replication groups.

### Global ARG occurrence

The clustering of ARGs in the domain network correlated with their predicted hosts (Supplemental Figure 3) and their geographical origin (Supplemental Figure 4). RrnaAD and Aminoglyc_resit were found in *Firmicutes* members, and lie in the mixed geographical cluster composed of rolling circle-replicating plasmids. ABC_tran, APH, and MFS_1 were found in *Acinetobacter* plasmids and lie in the European cluster of theta-replicating plasmids (Supplemental Figure 3 and 4). The association between ARGs and genera makes sense, as not all ARGs are beneficial in all bacterial genera.

Pentapeptide_4 (and the member gene qnrB19) was not associated with any group in the domain network. Accordingly, it was associated *Salmonella* plasmids which were not commonly predicted in the data. qnrB19-carrying plasmids seems to bear little resemblance to other plasmids, but they are found world-wide, especially in the Americas. The lack of plasmids similar to the qnrB19 plasmids together with the fact that *Salmonella*-plasmids was not normally found in sewage indicate that qnrB19 was not acquired in the wastewater environments. Nevertheless, their presence in the data

We saw no ResFinder hits on bacteriophages. This is in accordance with the finding that phage and plasmid domains are well separated in the network graph with the exception of the small overlap among domains for initiators of rolling circle replication.

The prevalence of macrolide resistance genes in sewage water across continents and species could indicate selective pressure for this trait in sewage water across the world. Though the genes msr(E) and mph(E) were almost only located in the European samples, but this could be a consequence of the composition of the microbiomes in these samples, as *Acinetobacter*, the carrier of the genes in sewage, was abundant in these samples. erm(B)-carrying plasmids were found worldwide, as were their host genera. It was recently indicated that methicillin resistance emerged in *Staphylococcus aureus* in hedgehogs as an adaptation to beta-lactam-producing dermatophyte^79^ highlights the coevolution among microbes in natural habitats as important sources of AMR emergence. A high percentage of antibiotics are not fully metabolized in humans, and macrolides have been detected in wastewater^80,81^, though wastewater treatment plants lower their abundance^81,82^.

It is strange that macrolide resistance selection apparently only affects *Firmicutes* and *Acinetobacter* species. It could of course be an issue of using machine learning models to predict genera; it should be kept in mind that the genera in this article are only predictions not experimentally verified. Nevertheless, it is notable that selective pressure towards macrolide resistance in sewage water seems to be a global phenomenon and it affects it affects widely different types of plasmids

## Conclusion

We found that the sewage water plasmidome was functionally distinct from sewage phages. There were a number combinations of plasmid backbone genes that were more common than others. These groups showed preferential co-occurrence with selected bacterial genera, with signals of geographical association to the sewage source, for example the higher larger amount of bacteriophages and viruses in Asian samples. Additionally, macrolide resistance-conferring plasmids were the most commonly found in the plasmid dataset. While msr(E) and mph(E) is mostly limited to *Acinetobacter* and European samples, erm(B) appeared on all continents, throughout a number of genera including important pathogens.

Macrolides are important antibiotics and their emergence in sewage water plasmidome could be a great threat to global health. Why macrolide resistance genes are the most prevalent is not evident, but it could be a consequence of the persistence of macrolides in wastewater. Another factor is the species composition of sewage water bacterial community. The common sewage water genus *Acinetobacter* carried a very large proportion of the macrolide resistance genes.

The prevalence of this genus in sewage water therefore to some extent explains the abundance of macrolide resistance in sewage plasmidomes. Though *Acinetobacter* is far from the most common human pathogen, the existence of a plasmid-carried reservoir of macrolide resistance is a cause for concern regardless.

The worldwide occurrence of ARGs is an important aspect of global health management. This study shows how sewage plasmidomes can be surveyed for the emergence of ARGs in the environment, and points to macrolide resistance as an important emergent gene in the global sewage plasmidome.

## Acknowledgements

The authors would like to thank Derya Aytan-Aktug for help with utilizing the tools PlasmidHostFinder and SourceFinder. The authors would also like to thank the two reviewers for their valuable comments and suggestions. This work was mainly funded by the Novo Nordisk Foundation Grant NNF16OC0021856 to FMA, and partially supported by funding from the European Union’s Horizon 2020 Research and Innovation programme under grant agreement No 773830: One Health European Joint Programme; FULL-FORCE project to FMA and SO.

